# BetaBuddy: An end-to-end computer vision pipeline for the automated analysis of insulin secreting β-cells

**DOI:** 10.1101/2023.04.06.535890

**Authors:** Anne M. Alsup, Kelli Fowlds, Michael Cho, Jacob M. Luber

**Author notes:** Corresponding Author (MC), (JML). These authors contributed equally to this work (Anne M. Alsup, Kelli Fowlds).

## Abstract

Insulin secretion from pancreatic β-cells is integral in maintaining the delicate equilibrium of blood glucose levels. Calcium is known to be a key regulator and triggers the release of insulin. This sub-cellular process can be monitored and tracked through live-cell imaging and subsequent cell segmentation, registration, tracking, and analysis of the calcium level in each cell. Current methods of analysis typically require the manual outlining of β-cells, involve multiple software packages, and necessitate multiple researchers - all of which tend to introduce biases. Utilizing deep learning algorithms, we have therefore created a pipeline to automatically segment and track thousands of cells, which greatly reduces the time required to gather and analyze a large number of sub-cellular images and improve accuracy. Tracking cells over a time-series image stack also allows researchers to isolate specific calcium spiking patterns and spatially identify those of interest, creating an efficient and user-friendly analysis tool. Using our automated pipeline, a previous dataset used to evaluate changes in calcium spiking activity in β-cells post-electric field stimulation was reanalyzed. Changes in spiking activity were found to be underestimated previously with manual segmentation. Moreover, the machine learning pipeline provides a powerful and rapid computational approach to examine, for example, how calcium signaling is regulated by intracellular interactions in a cluster of β-cells.

## Introduction

The Islets of Langherns are groups of endocrine cells that reside in the pancreas. They contain many cell types that significantly assist in glycemic regulation. The percent composition of cell types remains to be agreed upon, and varying methods have yielded different assessments [1]. Islets are typically comprised of approximately 50-70% β-cells [2–5] and 16-30% α-cells [2,5,6], with δ-, γ-, and ε-cells encompassing the remaining portion. There is an estimated 3.2 to 14.8 million islets in the human body, and they have been a major source of study due to their roles in hormone secretion and subsequent correlation to Diabetes Mellitus [7], which is a disorder that derives from the dysregulation of blood glucose levels. Type I Diabetes, specifically, is an autoimmune disorder characterized by the decreased secretion of insulin due to the destruction of β-cells. Complications from diabetes include micro- and macrovascular disorders such as thrombosis, atherosclerosis, and declines in cognitive function [8]. Two ongoing challenges to the treatment and/or cure of Type I Diabetes involve the reduced population of β-cells and the autoimmune attack within the body [9]. The main function of β-cells is the production and secretion of insulin. The release of this hormone is triggered by an increase in ATP [10] as the blood becomes oversaturated with glucose. As intense research efforts continue to explore this disease, many of the mechanisms behind insulin secretion are yet to be fully understood.

Insulin hormone dynamics is strongly correlated and regulated by calcium signaling. Glucose plays a large role in the stimulus of Ca^2+^ movement in the mitochondria, endoplasmic reticulum, nucleus, cytosol, and other areas of the cell and increases the level of cytosolic Ca^2+^ [11]. Additionally, an influx of Ca^2+^ through voltage-gated channels impacts insulin exocytosis and secretion. Insulin secretion patterns are typically studied by following the movement of Ca^2+^ oscillations (or spikes) in β-cells [12]. For example, membrane-permeable and high affinity calcium-sensitive fluorophores can be loaded into the cells. Upon binding free Ca^2+^ ion, fluorescent signals are generated and emitted when excited by an appropriate external light source. This process has been routinely captured in real-time subcellular imaging and allowed the investigator to track intracellular calcium concentration. While the Ca^2+^ dynamics in individual cells has been monitored and tracked [13], challenges remain to examine the Ca^2+^ responses in each cell, especially in cluster of cells that are more physiologically representative of islets. There is still much unknown regarding the effects of different stimuli on cell spiking activity, changes due to cell-cell interactions, and how insulin secretion fluctuates based on temporal or spatial variables. Additionally, measuring fluorescent intensities of individual cells that are clustered and/or overlap remains a daunting problem during image analysis. Efficient, accurate, and automated computational pipelines to determine potential cell-to-cell communication through Ca^2+^, are expected to enhance the current understanding of endocrine cell behaviors.

Automated image analysis tools are continually being developed and made readily available to the investigators. Technological advancements have both decreased the time required to perform wet lab experiments and enabled researchers to obtain more sophisticated data. Open-source software such as ImageJ [14] and its updated architecture, Fiji [15], have been widely utilized for the evaluation of biological images. Fiji’s plugin, Trackmate [16], can further be used for the tracking of individual cells. Multiple other tracking procedures have been written in Python for specific research needs [17–19]. These tools are among many that can be combined with customized algorithms for enhanced biological research.

Segmentation is an important tool that allows researchers to computationally identify regions of interest (ROIs) and analyze them using different methodologies [14]. Manual segmentation has in the past been a standard protocol. However, it is time intensive and often subject to human errors and biases (Supplemental Fig 1) [20,21]. Typically, analysis of fluorescent images begins by outlining individual cells to measure fluorescent intensities that might fluctuate with time. With the need for batch analysis of imaging data becoming more prevalent, an automated analysis tool is required to efficiently identify individual cells through a time-series image stack and eventually predict and validate physiological responses (e.g., insulin secretion). Hand segmentation remains a strong measure of evaluation for overall training accuracy, but automated methods are quickly being established as the new primary method of imaging analysis [22].

**Figure 1.**
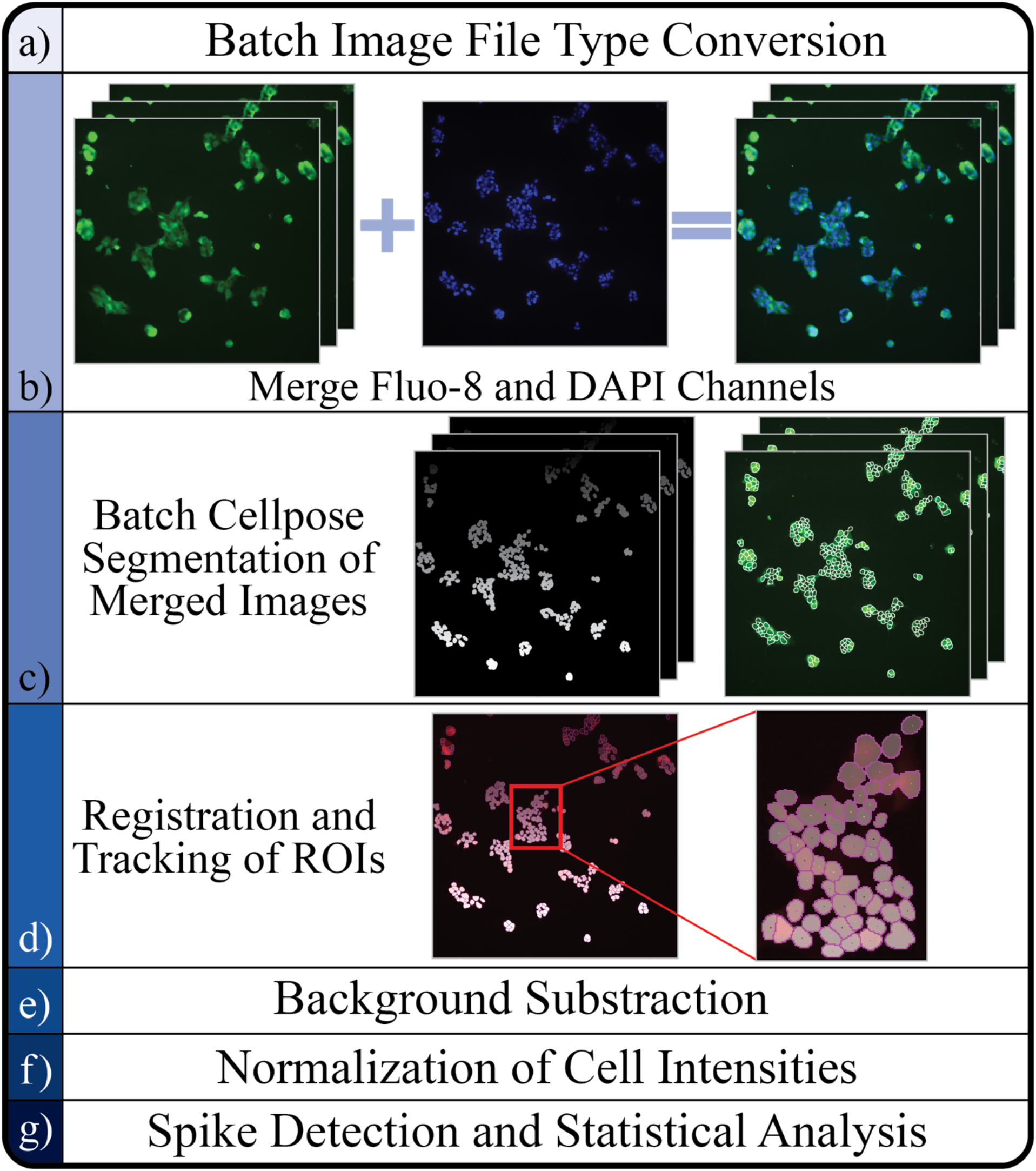
Pipeline Flow Chart. (a) Original Ca^2+^ microscopy images of β-cells (20x objective) converted to TIFF file from the Nikon-specific format. (b) Original Fluo-8 stained β-cells and DAPI stain of nuclei of β-cells showing only the nuclei of individual cells merged with both images in a separate channel, assisting in a more accurate segmentation. (c) ROI masks after complete image stack segmentation and overlayed predicted outlines from a sample frame after Cellpose segmentation (see below). (f) Visualization of tracks created after registration and tracking. This figure includes the original fluorescent image, mask image, ROI outlines, and predicted tracks cells followed throughout the imaging process. (e-g) Statistical analysis is incorporated to generated automated results, which can be compared with manually analyzed results (see Fig. 6 below).

In recent years, many tools have been developed to continue improving automated segmentation of cellular microscopy images including watershed algorithms, pre-processing techniques, deep learning methods such as Convolutional Neural Network architectures, and combinations of multiple techniques [23–29]. Before the wide use of deep learning, novel segmentation techniques involved differentiating cells from the background through thresholding, masking general cell outlines through complex mathematical operations, redefining cell shapes with the watershed algorithm, incorporation of active contours to identify rounded cell shapes, or a combination of multiple methods to create a more accurate segmentation of clustered cells [30–33]. While these previous techniques are still applicable and useful in conjunction with newer methods, the incorporation of deep neural networks has been able to overcome common issues such as heterogenous distribution of the fluorescent markers for watershed and uniform shapes making masking difficult. Deep learning-based methods are able to utilize large hand annotated data sets to accurately segment cells without relying on predefined shapes, distinct cell boundaries, or an evenly distributed cell culture with minimal clustering [34–38]. More novel deep learning networks, such as Cellpose, aim to create generalist models that increase flexibility of image analysis by training with a variety of cell types.

Cellpose [34] is a generalist, deep-learning algorithm designed for accurate segmentation of a wide variation of cell types. Its model is optimal for different types of biological research, as it provides segmentation without the need for retraining [34]. The algorithm is user-friendly and customizable for specific experimental needs, and its newest updates offer human-in-the-loop pipelines and methods for retraining [39]. Since its inception, several studies have already utilized its model. Saad et al. [40] developed a procedure using Cellpose and Fiji to characterize ice crystals, and Hoeren et al. [41] adapted Cellpose into their pipeline analyzing cells adhering to PMP-ECMO membranes as an alternative segmentation process to Fiji. Reinbigler et al. [42] modified the model’s base framework and used it in conjunction with QuPath [43] to create a pipeline for the segmentation of Hematoxylin-Eosin (HE) stained histopathological whole slide images of myofibers. Waisman et al. [44] similarly used Cellpose for myofiber segmentation, outputting the images to ImageJ for ROI analysis. Fisch et al. [45] incorporated StarDist [35] for nuclear segmentation and Cellpose for nuclear segmentation in their automated workflow for the analysis of host-pathogen interactions. Our aim in this study seeks to develop and apply computational methods to obtain a faster, more accurate and in-depth analysis of calcium signaling in β-cells.

## Materials and Methods

Our primary goal is to create an automated system that outlines cell boundaries (segmentation) and track individual β-cells over a sequence of different time points (Fig 1). The system includes and performs statistical analyses on the calcium intensity patterns from individual cells. The completed pipeline can be used through a Jupyter Notebook and all package dependencies are described in the GitHub repository. Referred to as the BetaBuddy GitHub, it also contains installation and usage instructions for inexperienced programmers to easily follow and create a more accessible tool for all researchers to implement in their experimental procedures. The automated pipeline is expected to improve the accuracy of data and decrease the overall time necessary for future studies. The entire procedure was run using an NVIDIA DGX A100 High Memory GPU Supercomputer, containing 8 NVIDIA A100 high memory graphics cards with 80GB of GPU RAM each for a total of 640GB of GPU RAM for training large Deep Bayesian model architectures. The system also has 2 TB of HBM2 RAM and 128 AMD EPYC CPU Cores, and it connects to the internet via 10 GB/S connection. The process from image conversion, segmentation, tracking, and finally statistical analysis was successfully performed with 1 GPU, 12 CPUs, and 32GB of memory allocated.

### Calcium imaging and deep learning-based segmentation

The images used in our automated pipeline were sourced from previous experimental images that monitored and recorded Ca^2+^ dynamics in real-time [46]. Mouse derived βTC-6 insulinoma cells were cultured and allowed to grow for several days. A calcium indicator dye (Fluo-8) was used to measure changes in intracellular calcium levels. The cells were stained with 0.8 μM Fluo-8, and 1 drop/mL Nucblue to visualize the nuclei, for 30 min at 37°C and loaded onto a custom-designed electric field exposure chamber [46]. Non-invasive electric field stimulation (EFS; 15 min exposure) was chosen to manipulate the voltage-gated calcium channels and therefore modulate intracellular Ca^2+^ dynamics. A Nikon microscope was used to image the cells at 5 s intervals over a period of 2 min before and after EFS (Figs. 1a and 1b). This method provided a baseline and changes in the intracellular Ca^2+^ levels were measured. Data for such experiments consisted of stacked 2048×2048 pixel image sets of 25 frames before and after EFS. In total, 56 image stacks were analyzed. The Fluo-8 fluorescent image data sets consisted of pre- and post-exposure images of non-invasive 1 to 3 V/cm EFS. The analysis pipeline was used to analyze approximately 9-10 image sets for each experimental condition. Pre-exposure cells served as their own control.

Once the cells are imaged, multiple single or stacked images can be imported directly to the script and begin segmentation after format conversion. The Java library, Bio-Formats, is first used to convert the Nikon-specific file types to TIFF for ease of use in Linux ImageJ macro functions. The stacked files are then merged with an image of only the nuclei (labeled with DAPI) from the first frame of the exposure period (Fig. 1b). It was determined that adding a DAPI channel greatly improved the segmentation process. Due to the clustering and static, non-migratory nature of β-cells, one DAPI image was determined sufficient in assisting segmentation for all frames.

A validated deep learning-based generalist cellular segmentation algorithm, Cellpose, is used to automatically outline the boundary of each cell and create an ROI with user-defined parameters [34]. Users may specify if their images contain a DAPI channel, which channel contains the entire cell, the Cellpose model for segmentation, and thresholds. These parameters can be tested on one image to ensure quality segmentation. The generalist algorithm has a better performance with the addition of a DAPI channel due to the clustering nature of β-cells and difficulty differentiating neighboring cells with only one calcium indicator dye. After segmentation, images defining the ROI of every cell are saved (Fig. 1c). ‘Mask’ images differentiate individual cells by assigning every ROI a different grayscale color (Fig. 1c). The unique color can be used as a label during cell registration and tracking.

The separated mask images are sorted into an image stack and merged with the original image with only Fluo-8. Merging is performed by sorting each image stack into a different color channel. This format will allow for the Fiji plugin, TrackMate, to begin cell registration by identifying the ROIs labeled in the channel containing the mask images [47]. The cell track detector analyzes the mask channel and registers every uniquely colored shape as an individual cell. Next, each cell is linked across multiple frames, creating a “track” (Fig. 1d). Lastly, fluorescent intensity for each frame in a cell’s track is obtained by referencing the first channel, which holds the original fluorescent image. The intensity values are averaged from within the cell’s complementary ROI. This process is completely automated using the ImageJ’s macro and python scripting features. This step will create an XML file that can be loaded into TrackMate and create a visualization of the tracks produced. Users are encouraged to check this visual before continuing to ensure the analysis is accurate. Lastly, a CSV file is saved containing fluorescent intensity values, area, and spatial coordinate positions by grouping each cell into a “Track ID” over time.

### Pipeline customization

BetaBuddy was designed as a customizable tool that can be easily adapted to the specific needs of each researcher. For this experiment, two primary modifications were made to the pipeline to assist in its data analysis. First, to analyze control vs. experimental data, we stitched the pre-exposure and post-exposure image sets together. Secondly, to account for the generalist algorithm not properly identifying low-intensity ROIs, we created a composite image of the maximum intensity value for every pixel, ensuring more definable cell boundaries for all active cells (Supplemental Fig 2). A composite image is created for the control and experimental image sets separately. Both composites are segmented, and the masks are overlayed on all frames in their respective image stacks. A flowchart for this pipeline can be seen in Supplementary Figure 2.

**Figure 2.**
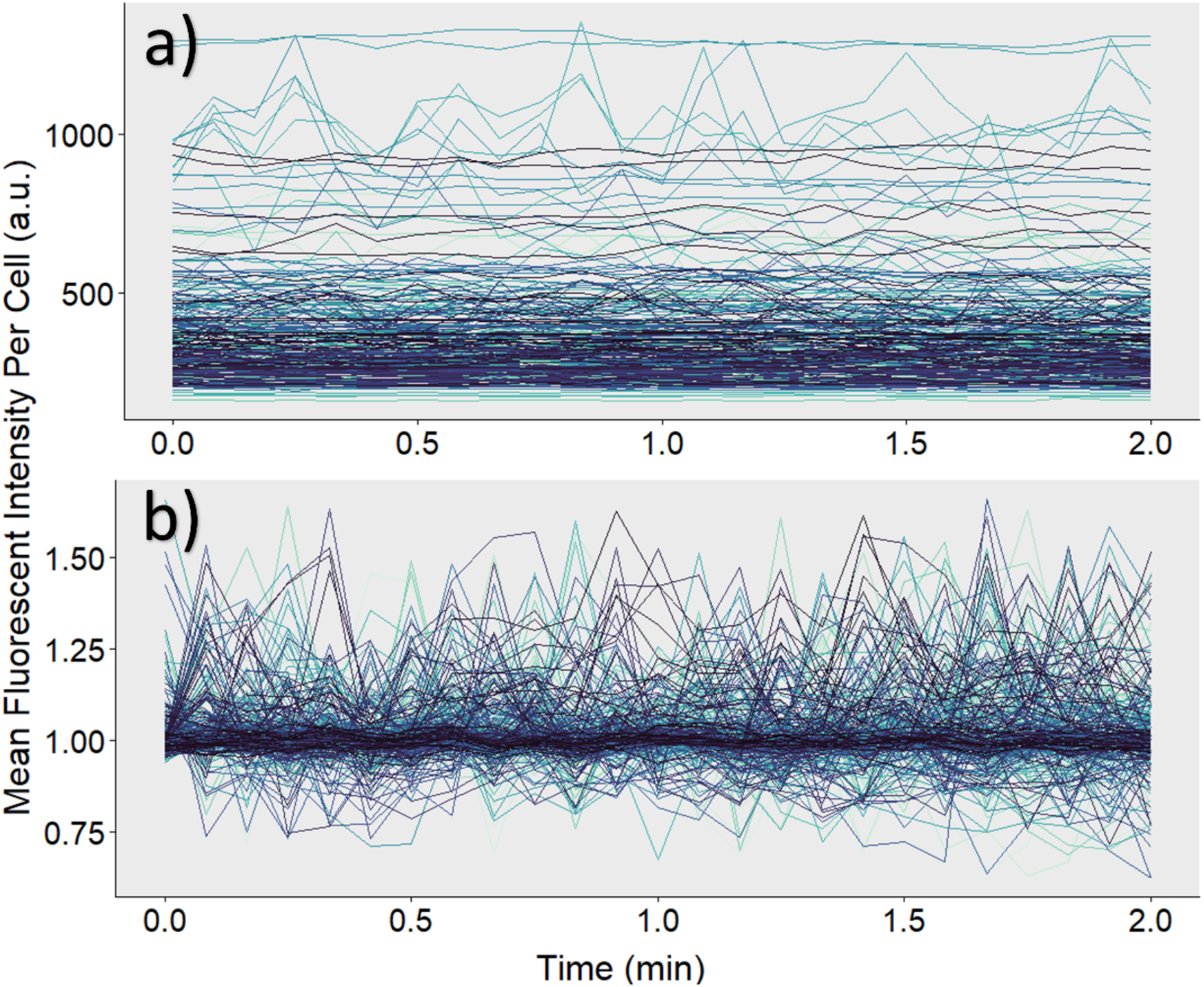
Intensity of Experimental Data. a) Each line represents an individual cell and its fluorescent intensity tracked over a period of 2 min. The same sample was treated with a 1 V/cm electric field for 15 min. b) Raw intensity values undergo background subtraction and normalization for a more accurate depiction of calcium activity.

### Background subtraction

Initially, the raw data CSV created after registration and tracking are very saturated with external background noise (Fig. 2a). Fluorescent images typically have background noise associated with excessive dye or thermal electronic fluctuation. Therefore, raw data should undergo a background subtraction process. To obtain a well-distributed sample of background intensities, 100 pixel locations outside of the ROIs are randomly selected from the original image stack. The randomly selected pixel is compared to the mask image stack to ensure the pixel intensity has a value of 0, verifying it is not within an ROI. The pixel intensities across all frames are saved into a CSV file and subtracted from the raw intensity values for normalization and statistical analysis (Fig. 2b).

### Normalization

Evaluation of all acquired data is analyzed through a procedure written in R and called upon through a terminal command in the Jupyter Notebook. The working directory is set to the user’s current operating space, and the CSV files containing the raw tracking information and background points are automatically imported. First, the average of all 100 randomly sampled background pixels is calculated for each unique frame. This mean value is then merged into the cell tracking information to create a singular data set. The average background intensity for each time point is then subtracted from the mean intensity of each cell within that frame. Next, to establish individual baseline activity, mean intensities were normalized specific to each individual cell.

In keeping with the original experiment’s methodology, a change in fluorescent intensity >=10% was notated as spiking activity [46]. We kept this 10% threshold for the purpose of comparing the accuracy of either manual or automated data analysis of calcium dynamics in β-cells. To find a baseline for each cell, the number of data points until each specific Track ID reached a change of >= 10% from its first recorded intensity was calculated, and the mean of these first N points was divided from each background-subtracted intensity within the group. If a cell never reached an intensity level greater than 10% of its first value, then its baseline was determined using the mean intensity of all recorded data points for that cell.

### Spike determination

The purpose of this experiment was to repeat the analysis of spiking activity within the original data set that utilized hand segmentation methods. Our goal was to determine if there was any significant loss of data in the original methodology. For the automated analysis, we defined a calcium spike using the 10% threshold, as previously stated. The percent change in normalized intensity from its previous value was calculated for each βTC-6 cell. Based on the value changes, a spike was defined in two ways: either (1) the percent change from the data point immediately preceding any peak in intensity was >= 10%, or (2) the sum of the cumulative percent changes between two peaks was >= 10% (Fig. 3).

**Figure 3.**
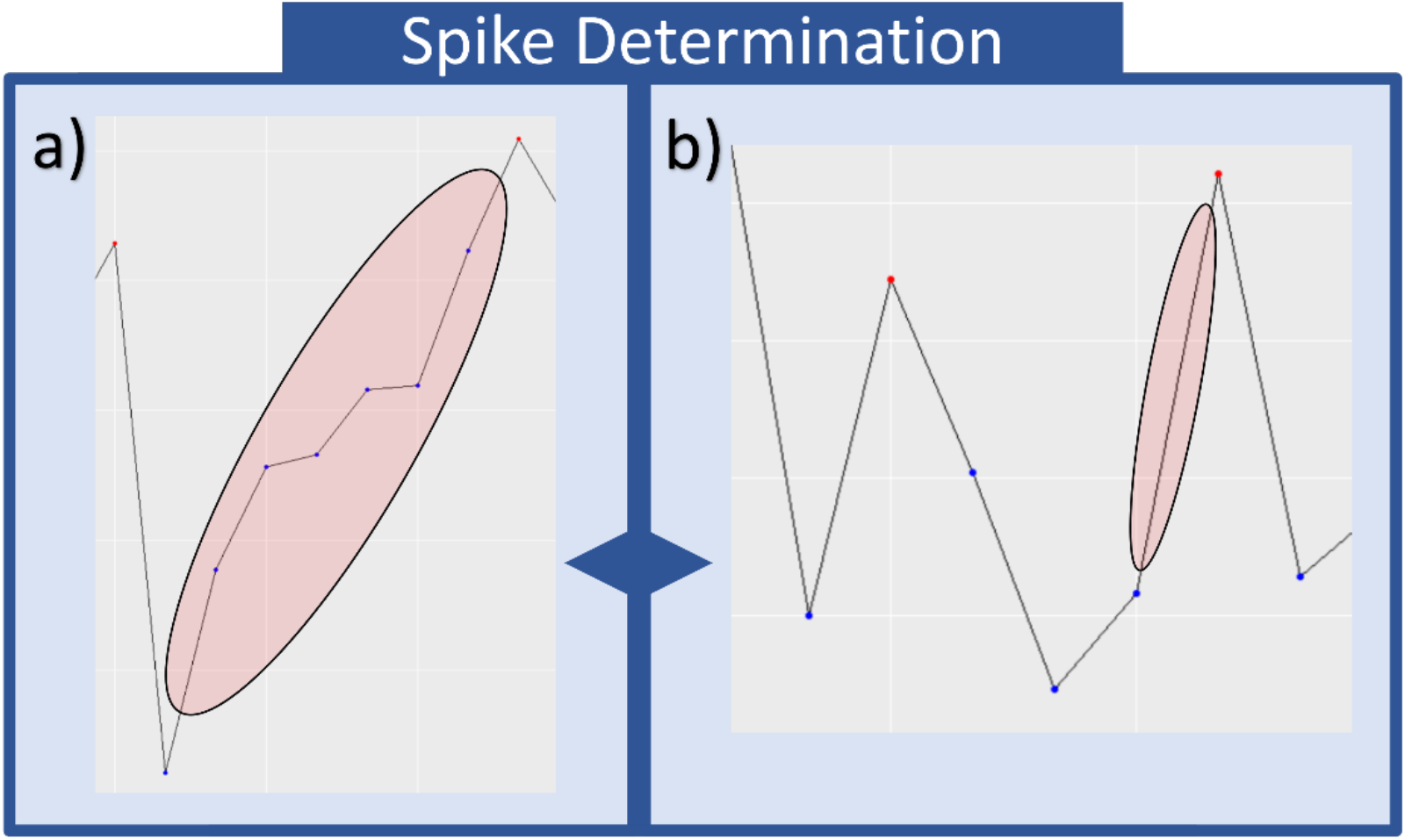
Calcium Spike Determination. Computational methods were used to automatically designate calcium spikes at a 10% threshold. This threshold was met through either (a) the sum of percent changes between two designated peaks or (b) a singular percent change of >= 10% immediately preceding a defined peak.

## Results

### Statistical analysis

Prior to statistical tests, outliers for individual trials were detected using a standard Z score. Track IDs with any intensity values greater than a Z score of 3.5, or less than -3.5 were automatically removed. Mean spiking activity was calculated as the total number of spikes recorded for each trial divided by total number of cells over time. A two-tailed paired *t* test was performed to compare changes in spiking activity between the pre- and post-EFS images of 1 to 3 V/cm. Figure 5 illustrates one experiment using 1 V/cm stimulation. Normalized calcium activity was plotted from -2 to 0 min (i.e., control before EFS was applied) and from 0 to 2 min following 15 min EFS. Calcium recording during 15 min EFS was not performed to minimize fluorophore degradation and/or bleaching.

**Figure 4.**
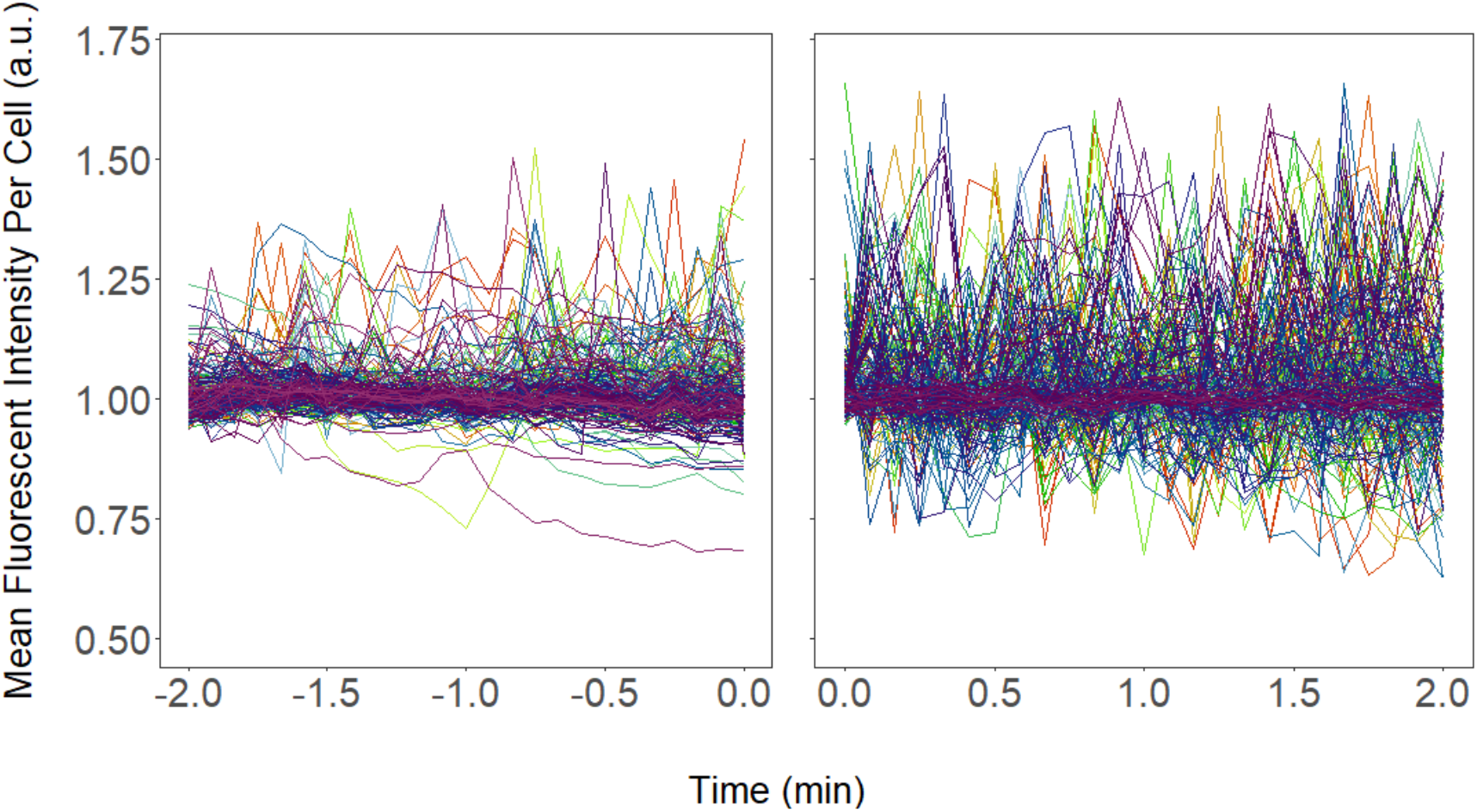
Signaling Changes Post EFS Stimulation. Variations in activity, including changes in the fluorescent intensity and spiking frequency, increased significantly following a 15 min exposure to 1 V/cm EFS. *** denotes p < 0.001.

**Figure 5.**
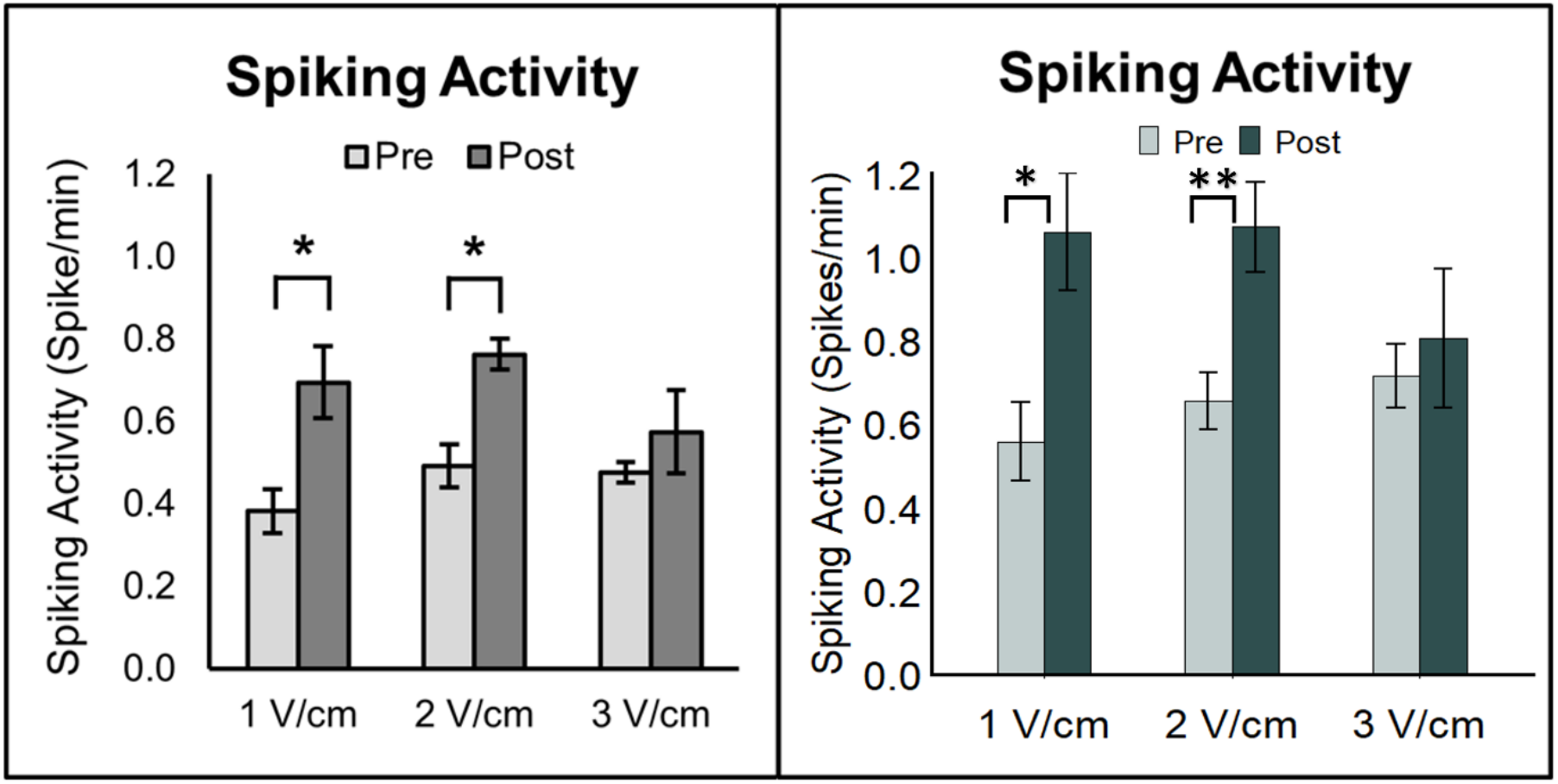
Comparison of Results. The number of calcium spikes determined by hand are shown in the left panel. Using the same images, the results from the automated analysis are shown in the right panel. * denotes p < 0.05 and ** p < 0.01.

Using our automated process, significant changes in the calcium signaling of the β-cells in response to EFS were determined and compared against published results. Applying EFS of 1 to 3 V/cm, the numbers of calcium spikes determined manually and using the automated pipeline were plotted and compared (Fig. 5). A clear trend is established that manually determined calcium spikes were underestimated by ∼ 40% in control cells without exposure to an EFS. In the EFS-treated cells (1 or 2 V/cm), the same trend is observed, suggesting that the calcium spike counting determined by hand is also underestimated. This finding is indicative of systematic errors that may have been caused by human biases. It is interesting to note that a larger EFS (3 V/cm) did not induce statistically significant changes in the calcium spiking determined either by hand or by automated pipeline. We speculated that, since the induced electrical potential depends on the cluster size, larger EFS causes non-negligible and perhaps irreversible changes in the cell membrane potential. Generally, automated analysis found greater overall spiking activity in the βTC-6 cells.

## Discussion

The application of electric field stimulation to βTC-6 cells was found to successfully increase calcium spiking activity. This indicates that EFS within 1 to 2 V/cm serves as a potentially beneficial treatment for pre-conditioning isolated islets prior to transplantation. While it remains to be further elucidated whether one-time treatment of islets with EFS is sufficient to pre-condition them, it does offer a non-biologic approach to regulate insulin secretion and/or trafficking. Development of the BetaBuddy pipeline therefore enables us to predict the optimal application of exogenous stimuli that can be validated experimentally and provides a pathway to fully utilize the machine learning-based discoveries in the β-cell physiology. Furthermore, results that were previously reported using manual segmentation (see Fig. 5) were compared with those determined by the BetaBuddy. Both sets of results were comparable, suggesting analysis by hand was able to quantify the calcium dynamics in β-cells but underestimated the calcium spiking rate. Automated segmentation thus appears to be a viable and more accurate method for obtaining reliable results without the tedious and laborious practices.

The output from this pipeline additionally offers considerably more information than would normally be available with standard segmentation practices. Previous analysis of this data only provided whole frame averages of fluorescent intensity changes. Specialized data for each individual cell at every experimental time point is available through the automated method. We are thus able to locate cells of interest and evaluate their spiking patterns based on specific temporal and spatial points. Questions regarding calcium spiking frequency and cell-to-cell interaction through calcium propagation are now feasible for detailed analysis. This is perhaps one of the major advantages that were made possible by the development of automated segmentation and analysis. For example, intercellular communication often relies on calcium waves through gap junctions [48–50].Since EFS we used in this work does not penetrate the cell, it may be hypothesized that the outer cells in a cluster of cells are likely affected by the EFS and generate calcium activities first. Through calcium intercellular communication, the inner cells in the cluster are affected at a later time. However, the cluster is collectively activated by EFS that leads to an increase in insulin production. Experiments are underway to validate this hypothesis and the results will again be used to train the model to recognize the calcium-dependent cell-to-cell communication that may be relevant to the islet physiology.

### Future work

Future work will include retraining the generalist segmentation pipeline, Cellpose, with a large set of β-cell images in order to create a specialized deep-learning model. Current segmentation with the generalist training is very useful and can give detailed insights on the image set, but it is not entirely accurate. Many cells become split into multiple ROIs, neighboring cells are merged into one mask, or cells will merge or split for one frame creating a broken track that will be misrepresented during normalization. We are theorizing that retraining with a massive dataset of only β-cell images and similarly clustering cells will give our pipeline the ability to create more accurate ROIs and subsequent individualized cell data for culturing condition and cell interaction studies.

Secondly, we plan to continue expanding the accessibility of our pipeline. A cleaner, more interactive user interface within the Jupyter Notebook is being developed along with an Anaconda package for researchers to easily create an environment with all the necessary packages pre-installed. Additionally, we are developing more user-friendly options for running different data analyses. This includes options for various statistical tests depending on researcher preference and the types of biological questions being asked. With these new features, users will be able to run desired tests within the pipeline along with having access to both the raw and normalized data.

This specialized data offers many resources for analyzing calcium signaling in new ways. Cell-specific spatial and fluorescent intensity data can be utilized, for example, to evaluate varying levels of insulin secretion in given locations (Fig 6). Researchers can use these methods to not only look at current spiking patterns, but also begin to analyze how calcium signaling is affected by intercellular interactions.

**Fig 6.**
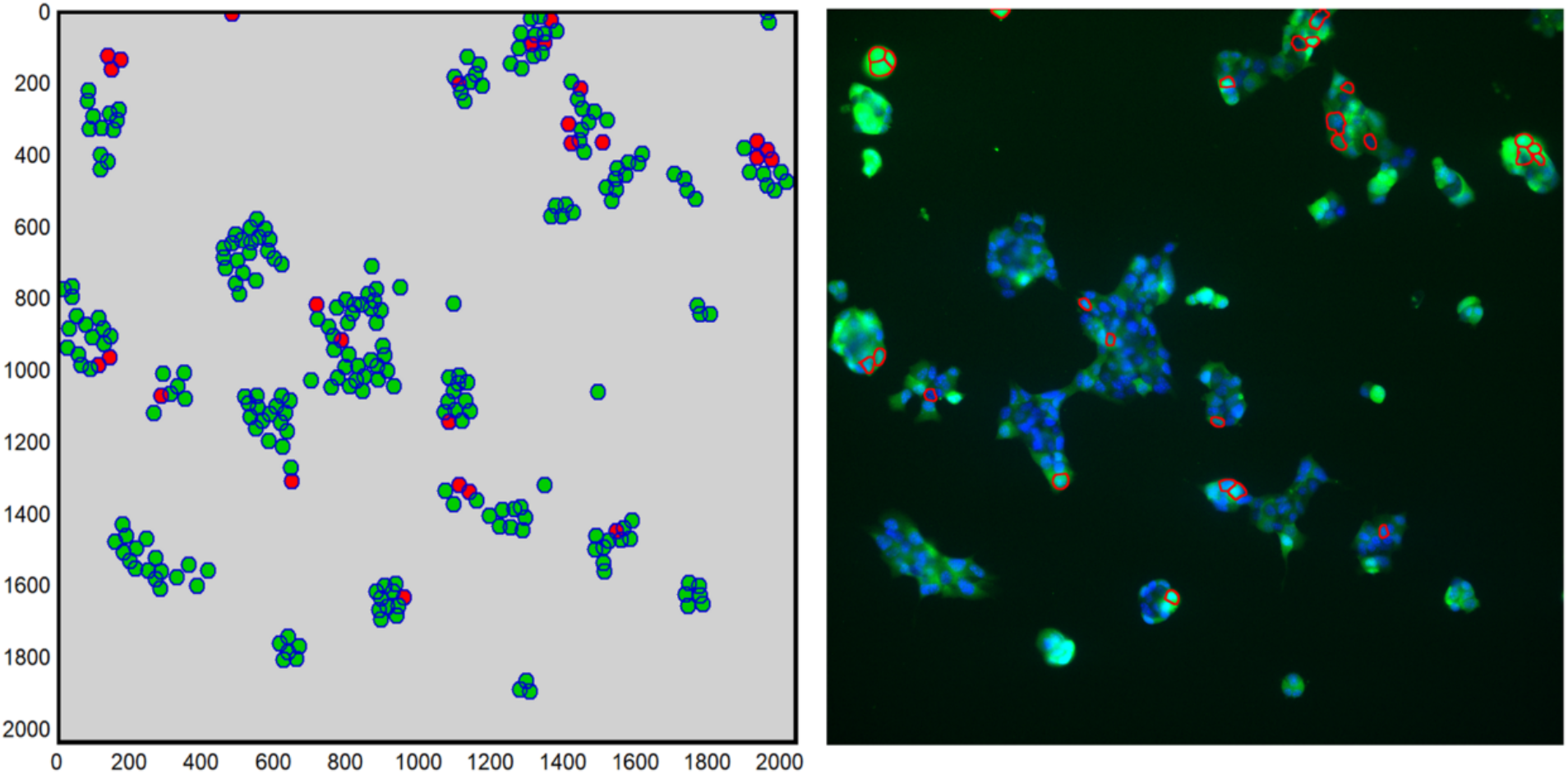
Spatial Visualization. a) Cartesian coordinates were used to visualize the locations of spiking cells. Only cells currently spiking are outlined at a given time. The cells highlighted in red color, for example, exhibit calcium spiking activity at t = 25 s. The visual analysis does not correlate to the actual size of cell clusters. b) The generated spatial data can be used to trace back and identify cells of interest in the original experimental images.

## Conclusions

Our current pipeline successfully isolates β-cells, tracks calcium signaling patterns over time, and produces preliminary data values that can be used in future statistical analyses. The automated segmentation additionally reduces biases, decreases time required for analysis compared with manual segmentation, and provides more individualized data. With this information, we are able to pinpoint individual cells with the most biologically significant spiking patterns. Additionally, individual calcium activities can be monitored over time, offering critical spatial and temporal information.

BetaBuddy has a multitude of benefits for potential research. Its customizable platform allows it to be adapted for use with multiple cell types. Its base structure has already been utilized for the automated analysis of mouse brain endothelial cells (MBECs) and human mesenchymal stem cells (hMSCs). Additionally, the pipeline is designed in a way that researchers from diverse background can use regardless of past coding experience.

Information obtained from this data may greatly assist in treatments for Type I Diabetes. Understanding the complexities of calcium signaling within pancreatic β-cells can provide guidance to better strategize and apply stem cell-derived islet transplants. There is still much to learn regarding the most effective forms of islet transplants, including clinically efficacious dosage and cell composition of stem cell-derived islets [9]. In the future, our algorithm has the potential to be used to predict viable cells for islet transplantation. Moreover, these methods can ultimately be used for the evaluation of invasive and non-invasive treatment options. Our automated analysis can potentially decrease evaluation times for multiple treatment alternatives, and even serve as a drug repurposing tool. This pipeline has the capability to lower the risk of failed transplants and will further provide the field with a valuable computational resource to significantly improve current treatment plans.

## Acknowledgements

We would like to thank Dr. Caleb Liebman for his dedicated work culturing insulinoma (βTC-6) mouse cells and providing the calcium images.

This work would not have been possible without the support of the NIH Predoctoral Training Grant (HL134613; PI, Kytai Nguyen), the University of Texas System Rising STARs Award (J.M.L), the CPRIT First Time Faculty Award (J.M.L), and the Alfred R. and Janet H. Potvin Endowment (M.C).

## Code and Data Availability

Directions for downloading the BetaBuddy pipeline and code used to perform this analysis can be found at the following public GitHub:

https://github.com/jacobluber/BetaBuddy

## Supplemental Figures

**Supplemental Figure 1.**
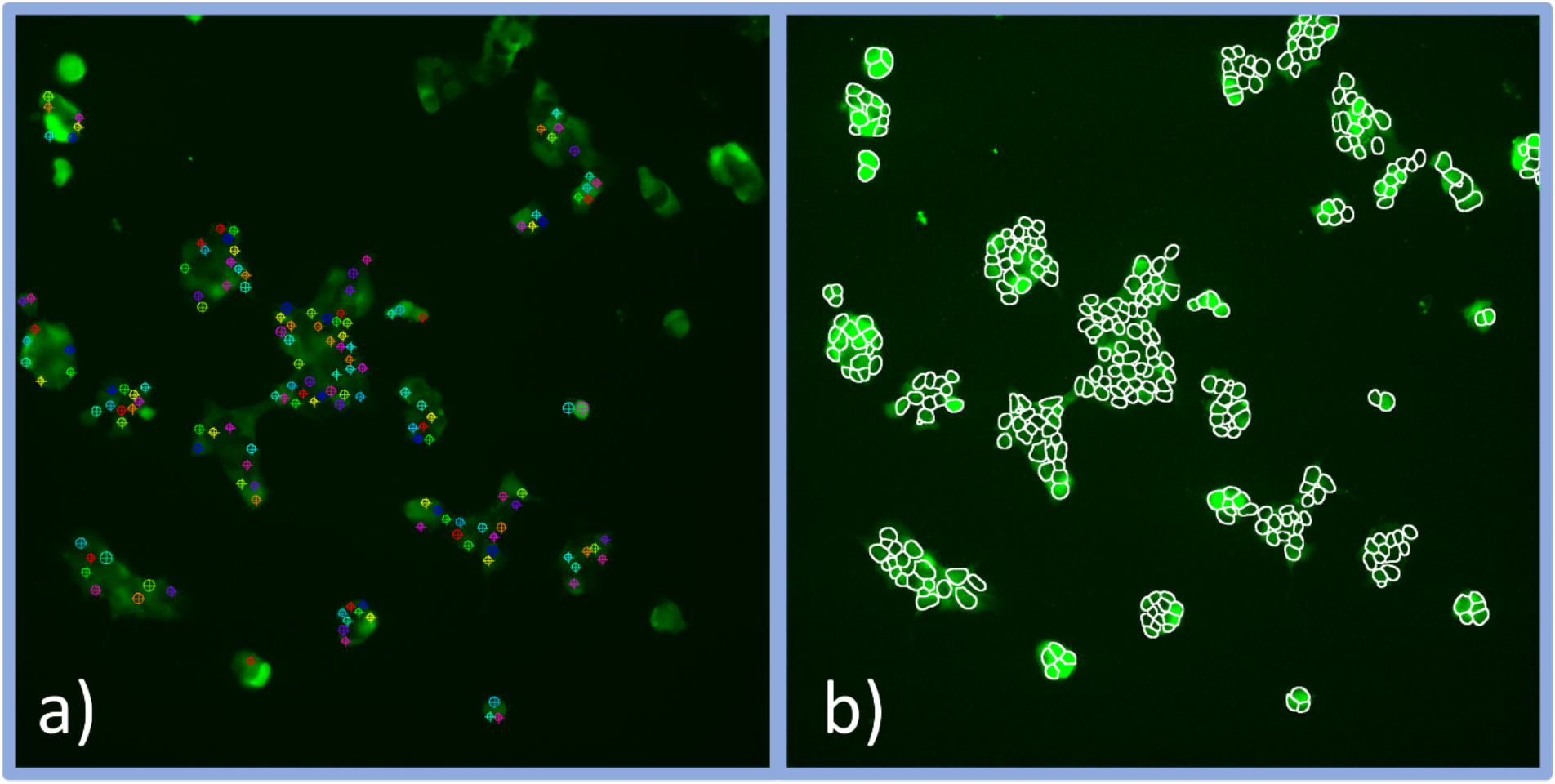
Manual vs. Automated Segmentation. Deep learning algorithms have been found to greatly improve segmentation accuracy. (a) Previous hand segmentation methods often undersegmented the total ROIs present, often ignoring highly clustered areas. (b) Our automated system comprising merging DAPI with the targeted fluorescein channel, segmentation, and subsequent ROI tracking was able to consistently identify more cells at a higher accuracy and track localized signals.

**Supplemental Figure 3.**
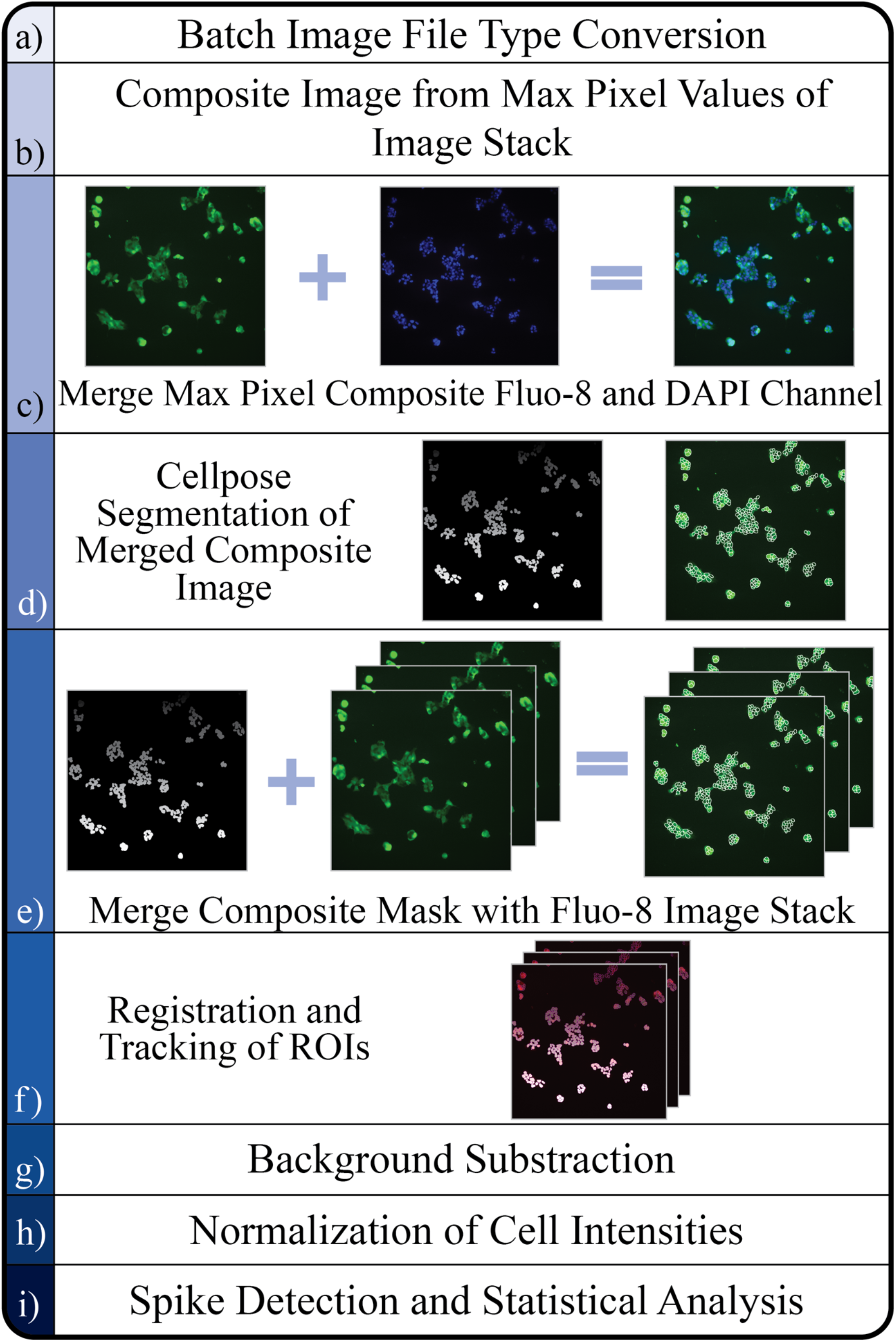
Adjusted BetaBuddy Pipeline Flow Chart. (a) Original Ca^2+^ microscopy images of β-cells (20x objective) converted to TIFF file from the Nikon-specific format. (b) A composite image is made by identifying the maximum value of each pixel location from the image stack. The resulting image is the brightest pixel out of all images in the stack, ensuring the states where each cell is fully fluorescing is in one image. This will allow for a more accurate segmentation without dropout due to non-fluorescing cells. (c) Composite Fluo-8 stained β-cell image and DAPI stain of nuclei of β-cells showing only the nuclei of individual cells merged with both images in a separate channel, assisting in a more accurate segmentation. (d) ROI mask after composite segmentation and overlayed predicted outlines from after Cellpose segmentation. (e) Mask of segmented composite image is merged with all images in the image stack. Due to this step only image stacks with exceptionally minimal movement should be used. (f) Visualization of tracks created after registration and tracking. This figure includes the original fluorescent image, mask image, ROI outlines, and predicted tracks cells followed throughout the imaging process. (e-g) Statistical analysis is incorporated to generated automated results, which can be compared with manually analyzed results.

**Supplemental Figure 3.**
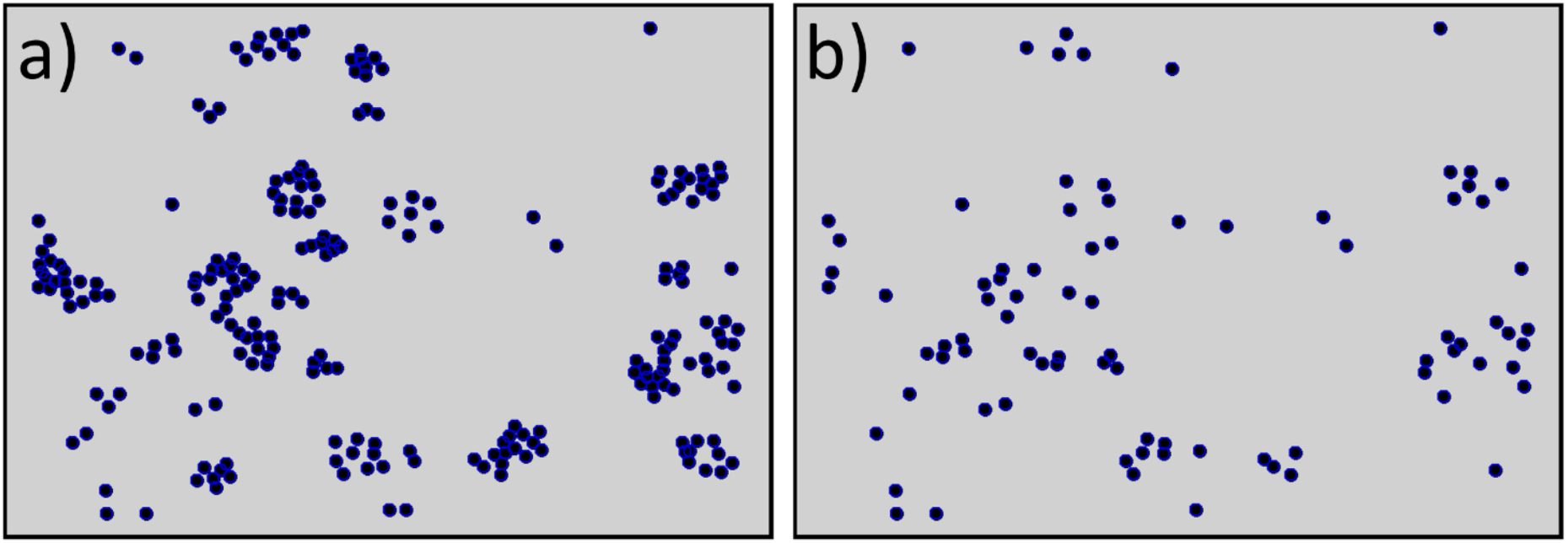
Previous Segmentation Loss. Initial registration and tracking was limited by subtle movements between imaging periods. The total number of unique ROIs (a) often decreased by ∼80% when accounting for only the registered cells that could be tracked between all frames (b). Customization of the BetaBuddy pipeline included merging the DAPI channel with the original Fluo-8 images and creating a composite image using the maximum intensity value for every pixel, allowing for optimum segmentation.

## References

1. Bonner-Weir S, Sullivan BA, Weir GC. Human Islet Morphology Revisited: Human and Rodent Islets Are Not So Different After All. J Histochem Cytochem. 2015 Aug 1;63(8):604–12.

2. Cabrera Over, Berman Dora M., Kenyon Norma S., Ricordi Camillo, Berggren Per-Olof, Caicedo Alejandro. The unique cytoarchitecture of human pancreatic islets has implications for islet cell function. Proc Natl Acad Sci. 2006 Feb 14;103(7):2334–9.

3. Butler AE, Janson J, Bonner-Weir S, Ritzel R, Rizza RA, Butler PC. β-Cell Deficit and Increased β-Cell Apoptosis in Humans With Type 2 Diabetes. Diabetes. 2003 Jan 1;52(1):102–10.

4. Pisania A, Weir GC, O’Neil JJ, Omer A, Tchipashvili V, Lei J, et al. Quantitative analysis of cell composition and purity of human pancreatic islet preparations. Lab Invest. 2010 Nov 1;90(11):1661–75.

5. Yoon KH, Ko SH, Cho JH, Lee JM, Ahn YB, Song KH, et al. Selective β-Cell Loss and α-Cell Expansion in Patients with Type 2 Diabetes Mellitus in Korea. J Clin Endocrinol Metab. 2003 May 1;88(5):2300–8.

6. Brissova M, Fowler MJ, Nicholson WE, Chu A, Hirshberg B, Harlan DM, et al. Assessment of Human Pancreatic Islet Architecture and Composition by Laser Scanning Confocal Microscopy. J Histochem Cytochem. 2005 Sep 1;53(9):1087–97.

7. Da Silva Xavier G. The Cells of the Islets of Langerhans. J Clin Med. 2018 Mar 12;7(3):54.

8. DiMeglio LA, Evans-Molina C, Oram RA. Type 1 diabetes. The Lancet. 2018 Jun 16;391(10138):2449–62.

9. Melton D. The promise of stem cell-derived islet replacement therapy. Diabetologia. 2021 May 1;64.

10. Liebman C, Loya S, Lawrence M, Bashoo N, Cho M. Stimulatory responses in α- and β-cells by near-infrared (810 nm) photobiomodulation. J Biophotonics. 2022 Mar 1;15(3):e202100257.

11. Klec C, Ziomek G, Pichler M, Malli R, Graier WF. Calcium signaling in ß-cell physiology and pathology: A revisit. Int J Mol Sci. 2019 Jan;20(24):6110.

12. Idevall-Hagren O, Tengholm A. Metabolic regulation of calcium signaling in beta cells. Adv Pancreat B Cell Funct Dysfunct. 2020 Jul 1;103:20–30.

13. Ashcroft FM, Rorsman P. KATP channels and islet hormone secretion: new insights and controversies. Nat Rev Endocrinol. 2013 Nov 1;9(11):660–9.

14. Schroeder AB, Dobson ETA, Rueden CT, Tomancak P, Jug F, Eliceiri KW. The ImageJ ecosystem: Open-source software for image visualization, processing, and analysis. Protein Sci. 2021 Jan 1;30(1):234–49.

15. Schindelin J, Arganda-carreras I, Frise E, Kaynig V, Longair M, Pietzsch T, et al. Fiji: an open-source platform for biological-image analysis. Nat Methods. 2012 Jul;9(7):676–82.

16. Tinevez JY, Perry N, Schindelin J, Hoopes GM, Reynolds GD, Laplantine E, et al. TrackMate: An open and extensible platform for single-particle tracking. Image Process Biol. 2017 Feb 15;115:80–90.

17. Schwanbeck J, Oehmig I, Dretzke J, Zautner AE, Groß U, Bohne W. YSMR: a video tracking and analysis program for bacterial motility. BMC Bioinformatics. 2020 Apr 29;21(1):166.

18. Kok RNU, Hebert L, Huelsz-Prince G, Goos YJ, Zheng X, Bozek K, et al. OrganoidTracker: Efficient cell tracking using machine learning and manual error correction. PLOS ONE. 2020 Oct 22;15(10):e0240802.

19. Mosby LS, Polin M, Köster DV. A Python based automated tracking routine for myosin II filaments. J Phys Appl Phys. 2020 May;53(30):304002.

20. Rizwan I Haque I, Neubert J. Deep learning approaches to biomedical image segmentation. Inform Med Unlocked. 2020 Jan 1;18:100297.

21. Laukamp KR, Pennig L, Thiele F, Reimer R, Görtz L, Shakirin G, et al. Automated Meningioma Segmentation in Multiparametric MRI. Clin Neuroradiol. 2021 Jun 1;31(2):357–66.

22. Voigt SP, Ravikumar K, Basu B, Kalidindi SR. Automated Image Processing Workflow for Morphological Analysis of Fluorescence Microscopy Cell Images. JOM. 2021 Aug 1;73(8):2356–65.

23. Gamarra M, Zurek E, Escalante HJ, Hurtado L, San-Juan-Vergara H. Split and merge watershed: A two-step method for cell segmentation in fluorescence microscopy images. Biomed Signal Process Control. 2019 Aug 1;53:101575.

24. Li G, Liu T, Nie J, Guo L, Chen J, Zhu J, et al. Segmentation of touching cell nuclei using gradient flow tracking. J Microsc. 2008;231(1):47–58.

25. Malpica N, de Solórzano CO, Vaquero JJ, Santos A, Vallcorba I, García-Sagredo JM, et al. Applying watershed algorithms to the segmentation of clustered nuclei. Cytometry. 1997 Aug 1;28(4):289–97.

26. Valen DAV, Kudo T, Lane KM, Macklin DN, Quach NT, DeFelice MM, et al. Deep Learning Automates the Quantitative Analysis of Individual Cells in Live-Cell Imaging Experiments. PLOS Comput Biol. 2016 Nov 4;12(11):e1005177.

27. Akram SU, Kannala J, Eklund L, Heikkilä J. Cell Segmentation Proposal Network for Microscopy Image Analysis. In: Carneiro G, Mateus D, Peter L, Bradley A, Tavares JMRS, Belagiannis V, et al., editors. Deep Learning and Data Labeling for Medical Applications. Cham: Springer International Publishing; 2016. p. 21–9.

28. S. E. Ahmed Raza, L. Cheung, D. Epstein, S. Pelengaris, M. Khan, N. M. Rajpoot. MIMO-Net: A multi-input multi-output convolutional neural network for cell segmentation in fluorescence microscopy images. In: 2017 IEEE 14th International Symposium on Biomedical Imaging (ISBI 2017). 2017. p. 337–40.

29. D. Eschweiler, T. V. Spina, R. C. Choudhury, E. Meyerowitz, A. Cunha, J. Stegmaier. CNN-Based Preprocessing to Optimize Watershed-Based Cell Segmentation in 3D Confocal Microscopy Images. In: 2019 IEEE 16th International Symposium on Biomedical Imaging (ISBI 2019). 2019. p. 223–7.

30. J. M. Sharif, M. F. Miswan, M. A. Ngadi, M. S. H. Salam, M. M. bin Abdul Jamil. Red blood cell segmentation using masking and watershed algorithm: A preliminary study. In: 2012 International Conference on Biomedical Engineering (ICoBE). 2012. p.258–62.

31. Koyuncu CF, Arslan S, Durmaz I, Cetin-Atalay R, Gunduz-Demir C. Smart Markers for Watershed-Based Cell Segmentation. PLOS ONE. 2012 Nov 12;7(11):e48664.

32. C. Zimmer, E. Labruyere, V. Meas-Yedid, N. Guillen, J.. -C. Olivo-Marin. Segmentation and tracking of migrating cells in videomicroscopy with parametric active contours: a tool for cell-based drug testing. IEEE Trans Med Imaging. 2002 Oct;21(10):1212–21.

33. I. Ersoy, K. Palaniappan. Multi-feature contour evolution for automatic live cell segmentation in time lapse imagery. In: 2008 30th Annual International Conference of the IEEE Engineering in Medicine and Biology Society. 2008. p. 371–4.

34. Stringer C, Wang T, Michaelos M, Pachitariu M. Cellpose: A generalist algorithm for cellular segmentation. Nat Methods. 2021 Jan 1;18(1):100–6.

35. Schmidt U, Weigert M, Broaddus C, Myers G. Cell Detection with Star-Convex Polygons. In: Frangi AF, Schnabel JA, Davatzikos C, Alberola-López C, Fichtinger G, editors. Medical Image Computing and Computer Assisted Intervention – MICCAI 2018. Cham: Springer International Publishing; 2018. p. 265–73.

36. Apthorpe N, Riordan A, Aguilar R, Homann J, Gu Y, Tank D, et al. Automatic Neuron Detection in Calcium Imaging Data Using Convolutional Networks. In: Advances in Neural Information Processing Systems [Internet]. Curran Associates, Inc.; 2016 [cited 2023 Mar 30]. Available from: https://proceedings.neurips.cc/paper/2016/hash/0771fc6f0f4b1d7d1bb73bbbe14e0e31-Abstract.html

37. Liu Z, Jin L, Chen J, Fang Q, Ablameyko S, Yin Z, et al. A survey on applications of deep learning in microscopy image analysis. Comput Biol Med. 2021 Jul 1;134:104523.

38. Xu J, Zhou D, Deng D, Li J, Chen C, Liao X, et al. Deep Learning in Cell Image Analysis. Intell Comput [Internet]. [cited 2023 Mar 30];2022. Available from: https://doi.org/10.34133/2022/9861263

39. Pachitariu M, Stringer C. Cellpose 2.0: how to train your own model. Nat Methods. 2022 Dec 1;19(12):1634–41.

40. Saad J, Fomich M, Día VP, Wang T. A novel automated protocol for ice crystal segmentation analysis using Cellpose and Fiji. Cryobiology. 2023 Jun 1;111:1–8.

41. Hoeren F, Görmez Z, Richter M, Troidl K. Deetect: A Deep Learning-Based Image Analysis Tool for Quantification of Adherent Cell Populations on Oxygenator Membranes after Extracorporeal Membrane Oxygenation Therapy. Biomolecules. 2022 Dec 3;12(12):1810.

42. Reinbigler M, Cosette J, Guesmia Z, Jimenez S, Fetita C, Brunet E, et al. Artificial intelligence workflow quantifying muscle features on Hematoxylin–Eosin stained sections reveals dystrophic phenotype amelioration upon treatment. Sci Rep. 2022 Nov 19;12:19913.

43. Bankhead P, Loughrey MB, Fernández JA, Dombrowski Y, McArt DG, Dunne PD, et al. QuPath: Open source software for digital pathology image analysis. Sci Rep. 2017 Dec 4;7(1):16878.

44. Waisman A, Norris AM, Elías Costa M, Kopinke D. Automatic and unbiased segmentation and quantification of myofibers in skeletal muscle. Sci Rep. 2021 Jun 3;11:11793.

45. Fisch D, Evans R, Clough B, Byrne SK, Channell WM, Dockterman J, et al. HRMAn 2.0: Next-generation artificial intelligence–driven analysis for broad host–pathogen interactions. Cell Microbiol. 2021;23(7):e13349.

46. Liebman C, Vu TM, Phillips A, Chen B, Cho M. Altered β-cell calcium dynamics via electric field exposure. Ann Biomed Eng. 2021 Jan;49(1):106–14.

47. Ershov D, Phan MS, Pylvänäinen JW, Rigaud SU, Le Blanc L, Charles-Orszag A, et al. Bringing TrackMate into the era of machine-learning and deep-learning. bioRxiv. 2021 Jan 1;2021.09.03.458852.

48. Benninger RKP, Head WS, Zhang M, Satin LS, Piston DW. Gap junctions and other mechanisms of cell–cell communication regulate basal insulin secretion in the pancreatic islet. J Physiol. 2011 Nov 1;589(22):5453–66.

49. Evans WH, Martin PEM. Gap junctions: structure and function (Review). Mol Membr Biol. 2002 Jan 1;19(2):121–36.

50. Benninger RKP, Zhang M, Head WS, Satin LS, Piston DW. Gap Junction Coupling and Calcium Waves in the Pancreatic Islet. Biophys J. 2008 Dec 1;95(11):5048–61.

